# Coordinated Evolution of Influenza A Surface Proteins

**DOI:** 10.1101/008235

**Authors:** Alexey D. Neverov, Sergey Kryazhimskiy, Joshua B. Plotkin, Georgii A. Bazykin

## Abstract

Surface proteins hemagglutinin (HA) and neuraminidase (NA) of the human influenza A virus evolve under selection pressure to escape the human adaptive immune response and antiviral drug treatments. In addition to these external selection pressures, some mutations in HA are known to affect the adaptive landscape of NA, and vice versa, because these two proteins are physiologically interlinked. However, the extent to which evolution of one protein affects the evolution of the other is unknown. Here we develop a novel phylogenetic method for detecting the signatures of such genetic interactions between mutations in different genes – that is, intergene epistasis. Using this method, we show that influenza surface proteins evolve in a coordinated way, with substitutions in HA affecting substitutions in NA and vice versa, at many sites. Of particular interest is our finding that the oseltamivir-resistance mutations in NA in subtype H1N1 were likely facilitated by prior mutations in HA. Our results illustrate that the adaptive landscape of a viral protein is remarkably sensitive to its genomic context and, more generally, imply that the evolution of any single protein must be understood within the context of the entire evolving genome.

## Author summary

A single trait may be affected by several genes. But are the effects of different genes on a trait independent from each other, or do genes interact to determine the phenotype? The prevalence and type of such interactions is a topic of hot debate. They can be inferred from time-series sequencing data when a change in one gene is observed to facilitate a change in another gene. However, the situation is complicated when new combinations of genes are formed by processes such as recombination or reassortment. In such cases, deducing the time and order of the genetic changes is hard. Here, we devise a method to infer pairs of mutations that closely follow one another in presence of recombination. We apply it to evolution of two surface proteins of influenza A virus, hemagglutinin and neuraminidase, that are important targets of immune system and drugs; and show that mutations in one of these proteins are often permitted by prior mutations, or compensated by subsequent mutations, in the other. In particular, drug-resistance mutations in neuraminidase were made possible by prior mutation in hemagglutinin. Knowledge of such interactions is necessary to fully understand and, possibly, predict the evolution.

## Introduction

One of the central obstacles in controlling many pathogen-borne diseases is their exceptional ability to adapt through evolutionary changes [1]. Large population sizes and high mutation rates in many pathogens make them extremely effective at evolving to evade the immune system or resist drug treatments [2–6]. Our ability to prevent or even predict such escape mutations is hampered by limited knowledge of the effects of new mutations on pathogen fitness. This problem is made especially difficult because the effect of any particular mutation is often highly dependent on the genetic background in which it occurs, a phenomenon called epistasis [7–16].

Epistasis is particularly common among mutations that arise in response to strong selection pressures. For example, resistance mutations that arise under drug treatments often carry substantial fitness costs which are alleviated by secondary, compensatory, mutations [7,10, 14–16]. Likewise, mutations that facilitate immune escape are in several cases known to be epistatic with other, compensatory or permissive, mutations [17,18]. The surface proteins hemagglutinin (HA) and neuraminidase (NA) of the human influenza A virus evolve under strong selection pressures imposed by the human immune system and, possibly, antiviral drugs [4,19]. It is therefore expected that epistasis may play an important role in the evolution of these proteins. Several previous studies have found that epistasis within each of these proteins is widespread, so that mutations in a given protein are often beneficial only in the presence of mutations at other sites in the same protein [19–21].

Aside from intra-gene epistasis, we also might expect inter-gene epistasis, especially in the case of the HA and NA proteins of influenza viruses, which serve complementary physiological functions. HA facilitates the attachment of the virus to the cell surface, whereas NA catalyzes the separation of the ready-made virus particles from the cell. Thus, mutations that increase receptor-binding avidity of HA should promote mutations in NA that increase its cleavage activity [22,23] and vice versa [24,25]. HA and NA jointly determine sensitivity to neuraminidase inhibitors, with mutations in HA compensating for the reduction in binding affinity of NA caused by the inhibitors [26]. Other, as yet unknown, molecular interaction mechanisms may also lead to inter-gene epistasis. Indirect evidence also suggests that interactions between HA and NA may be strong; for example, reassortments giving rise to new combinations of HA and NA lead to a temporary increase in the rate of substitutions in these genes, likely due to accumulation of changes adjusting the genes to each other [27,28].

Here we present a method to detect signatures of inter-gene epistasis, and apply it to understand the evolutionary history of influenza surface proteins. The method we develop is an extension of techniques previously developed for detecting intra-gene epistasis [21,29]. The idea behind it is simple: epistasis will tend to induce temporal clustering of substitutions along the phylogeny of an adapting protein, with substitutions at one site followed rapidly by substitutions at another, interacting site. In the case of mutations within a single protein this idea can easily be developed into a rigorous statistical test, by quantifying the time that separates subsequent substitutions along the protein's phylogeny. All the sites within a single influenza protein share a common phylogenetic history: recombination events within an influenza virus RNA segment are exceedingly rare [30], and so sites that reside on the same segment of the viral genome are completely linked. However, influenza viruses undergo frequent reassortment events, so that sites residing on different segments typically have different genealogies – a complication that obscures the temporal order of substitutions occurring on different RNA segments. To account for this complication, here we develop a method for inferring the relative temporal order of substitutions at sites that have different evolutionary histories, and then use this information to detect temporal clustering of such substitutions in influenza viruses. We find that substitution rates at many sites in NA facilitate following substitutions in HA and vice versa, implying that inter-gene epistasis has shaped the molecular evolution of influenza viruses.

## Results

### Inferring reassortment events between HA and NA genes

We reconstructed individual phylogenetic trees for each of the two surface proteins HA and NA for the two major influenza subtypes circulating in humans, H3N2 and H1N1. Using software GIRAF [31], we identified taxa that descended from within-subtype reassortant ancestors and thus inferred the positions of reassortment events on the phylogenies of individual segment (see Materials and Methods for details). We inferred a total of 15 reassortment events between these two segments in subtype H3N2, and 5 events in subtype H1N1. We found that 847 out of 1,376 H3N2 isolates and 201 out of 745 H1N1 isolates are descendants of at least one reassortment event, which is consistent with previous findings [28,32,33]. To completely resolve incongruencies between individual segment phylogenies, we assumed that reassortments are the only source of true differences between the phylogenies of individual segments. This assumption imposes a constraint that phylogenies of different segments may differ by at most as many rooted subtree prune-regraft (rSPR) operations as there are reassortment events, and otherwise be identical. We reconstructed such “constrained” phylogenies of individual segments using previously inferred “unconstrained” individual segment phylogenies as templates (see Materials and Methods for details).

### Identifying pairs of sites involved in positive inter-gene epistasis

Acceleration of the rate of substitutions at one site (referred to as “trailing” site) following a genetic change at another site (referred to as “leading” site) indicates that mutations at the trailing site are more beneficial after a mutation occurs at the leading site, and thus indicate positive epistasis [21,29]. Here we are specifically interested in situations when leading and trailing sites are located in different genes and therefore have potentially different evolutionary histories. This fact complicates the inference of the temporal order of substitutions. Consider substitutions *i* and *ii* in the toy example presented in Figure 1A. While both of them obviously occurred on the line of descent of isolate *b*, it is not immediately clear whether substitution *i* in segment 1 occurred before or after substitution *ii* in segment 2. We therefore cannot say a priori whether substitution *ii* accelerated substitution *i*, or substitution *i* accelerated substitution *ii*, or there was no interaction between them at all.

**Figure 1.**
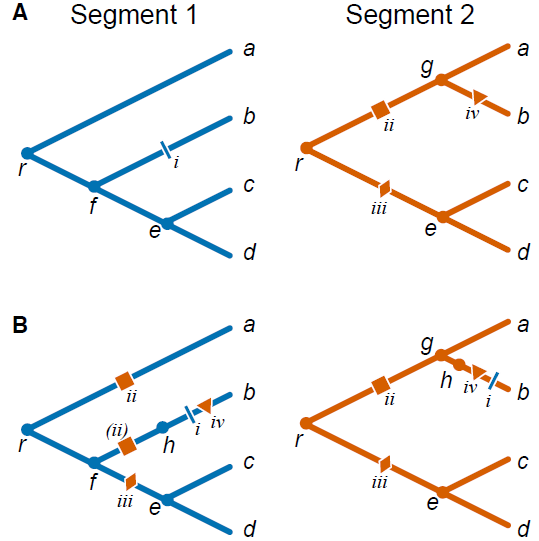
Mapping substitutions between segments in the presence of reassortments. (**A)** Individual toy phylogenies for segments 1 (left) and 2 (right) with respective substitutions. **(B)** Segment 1 phylogeny with segment 2 as the genetic background (left) and segment 2 phylogeny with segment 1 as the genetic background (right). *a–d,* leaf nodes; *e–g,* internal nodes; *h,* virtual node arising due to the reassortment event; *r,* root node; nodes corresponding to the two segments of the same isolate are denoted with the same letter. Each substitution is identified by a roman numeral and a unique symbol colored according to the segment in which it occurred. Substitution *ii* in segment 2 maps onto two branches of the segment 1 phylogeny, once as a regular substitution onto branch *ra* and once as a virtual substitution (denoted by parentheses) onto branch *fh*. Circles represent reassortment events.

To resolve such ambiguities, we estimate the temporal order of substitutions in different genes using the constrained phylogenies constructed above. Specifically, in order to study acceleration in substitution rates in one gene (referred to as the “foreground” gene) that follow substitutions in the other gene (referred to as the “background” gene), we map all non-synonymous substitution in the background gene onto the phylogeny of the foreground gene. Since the constrained phylogenies are topologically identical with the exception of a relatively small number of reassortment events, most branches of the background tree correspond to unique branches of the foreground tree. Most substitutions in the background gene are therefore unambiguously mapped onto branches of the foreground-gene phylogeny. In the toy example shown in Figure 1 branches *gb*, *ec*, and *ed* in the segment 2 phylogeny correspond to branches *fb*, *ec*, *ed* of the segment 1 phylogeny, respectively. Therefore, when considering segment 2 as the background gene, substitution *iv* unambiguously occurs on branch *fb* of the segment 1 phylogeny (Figure 1B).

Ambiguities in mapping background-gene substitutions onto the foreground-gene phylogeny arise at branches that precede and follow reassortment events, such as branches *rg*, *ga*, and *re* in the segment 2 phylogeny in Figure 1A. For instance, substitution *iii* in segment 2 could occur either on branch *rf* or on branch *fe* of the segment 1 phylogeny. We resolve such ambiguities by placing background substitutions onto the distal branch of the foreground phylogeny (e.g., in Figure 1, substitution *iii* is placed on branch *fe*). This choice minimizes the potential number of substitution pairs that contribute to our epistasis statistic (see below and Materials and Methods).

Finally, reassortment events themselves represent genetic changes in the background gene which may potentially elicit epistatic responses in the foreground gene. Indeed, when viewed as an event on the line of descent of a foreground-gene isolate, each reassortment event is a replacement of the genetic background gene, equivalent to multiple simultaneous substitutions that we call “virtual”. To account for the possibility that some of such virtual substitutions in the background gene lead to acceleration in substitution rates at foreground-gene sites, we mark each reassortment events by a “virtual” node on the foreground-gene phylogeny. All foreground-gene substitutions that occur on the respective branch are then placed after the virtual node. Here we make a simplifying assumption that reassortment events precede all substitutions on the respective branch (see Materials and Methods for details). Even though this assumption introduces an error in our inference of relative order of substitutions, this error is likely to be small because the fraction of braches involved in reassortment events is small. To illustrate this procedure, consider again the toy example in Figure 1. When considering segment 1 as the foreground gene, we posit that the reassortment (virtual node *h*) precedes substitution *i* on branch *fb* (segment 1) and substitution *iv* on branch *gb* (segment 2) which is also mapped onto branch *fb* of the segment 1 phylogeny (Figure 1B). This reassortment event replaces background segment 2 variant that carries no mutations (present at node *f*) with a segment 2 variant descendent from node *g* that has mutation *ii*. Thus, substitution *ii* is a virtual substitution in the background gene, and is placed on the virtual branch *fh*. Note that substitution *ii* is also mapped onto branch *ra* of segment 1 phylogeny.

Once all genetic changes in the background gene are mapped onto the foreground-gene phylogeny, we can use our previously developed method [21] for detecting acceleration in the rate of substitutions at sites in the foreground gene that follow substitutions in the background gene. To do so, we compute the epistasis statistic for each pair of sites *(i, j)* where the leading site *i* is in the background gene and the trailing site *j* is in the foreground gene (see Materials and Methods). The epistasis statistic tends to be large for those pairs of sites in which a non-synonymous substitution at the trailing site quickly follows a non-synonymous substitution at the leading site and for which such substitutions at the trailing site occur in multiple descendant lineages. We measure time between non-synonymous substitutions at these sites as the number of synonymous substitutions in the foreground gene that occur between them. As in our previous study [21], we exclude all substitutions at terminal branches because many such substitutions are likely to be deleterious.

Finally, to identify the pairs of sites with the epistasis statistic greater than expected by chance, we randomly reshuffle foreground-gene substitutions among branches of the foreground-gene phylogeny while keeping the mapped background-gene substitutions fixed. This procedure breaks all potential associations between background-and foreground-gene substitutions, while preserving the total number of substitutions at each site and on each branch of the phylogeny. It produces the null distributions of the epistasis statistic for all pair of sites simultaneously and allows us to estimate the false discovery rate (FDR) for any desired nominal *P*-value threshold [21].

### Prevalence of inter-gene epistasis in influenza surface proteins

We considered both HA and NA as foreground and background genes, for both subtypes. In all cases, we found a significantly higher number of site pairs with unexpectedly high values of the epistasis statistic in our data, implying abundant positive inter-gene epistasis (Table 1 and S3–S6, Figures 2–5 and S1). The observed number of epistatic pairs was significantly greater than expected for all considered nominal *P*-value thresholds below 0.05 in three of the four comparisons ((background N2, foreground H3), (background H3, foreground N2), and (background H1, foreground N1)); in the fourth comparison (background N1, foreground H1), it was significant for nominal p-values of 0.005 and below (Figures S1-S4). To form conservative lists of epistatically interacting pairs of sites, we chose the threshold nominal *P*-values that minimizes the FDR, while still retaining enough sites for the downstream analyses (Table 1, see Materials and Methods). At these thresholds, the number of epistatically interacting pairs of sites is about 5 times greater than expected by chance in three of the four comparisons ((background N2, foreground H3), (background H3, foreground N2), and (background H1, foreground N1)), and about 2.5 times greater than expected in the remaining comparison (background N1, foreground H1) (Table 1).

**Table 1.** Pairs of sites in HA and NA evolving under positive inter-gene epistasis.

| Subtype | H3N2 |  | H1N1 |  |
| --- | --- | --- | --- | --- |
| Number of sequences | 1,376 |  | 745 |  |
| Gene pair (background, foreground) | (N2,H3) | (H3,N2) | (N1,H1) | (H1,N1) |
| Foreground protein sites, total | 563 | 459 | 566 | 470 |
| Foreground protein sites, variable* | 173 | 147 | 130 | 122 |
| Total number of site pairs | 25,431 | 25,431 | 15,860 | 15,860 |
| $\tau$ used** | 62 | 50 | 62 | 52 |
| Nominal $P$-value threshold | $5 \times 10^{-4}$ | $5 \times 10^{-4}$ | 0.001 | 0.001 |
| Significant pairs (expected) | 5.92 | 4.78 | 5.76 | 5.18 |
| Significant pairs (observed) | 29 | 24 | 13 | 25 |
| FDR, % | 20 | 20 | 44 | 21 |
| Distinct leading sites | 23 | 21 | 13 | 25 |
| Distinct trailing sites | 13 | 11 | 5 | 6 |

\*Number of sites variable on internal branches
\*\*Value of the timescale parameter $\tau$ [21]

At these thresholds, between 11% and 19% of sites were involved in epistasis as leading, and between 4% and 8%, as trailing, depending on the considered pair of genes. For example, among the variable sites of H3, 8% (13/173) were involved in epistasis with N2 as trailing sites, and 12% (21/173), as leading sites (Table 1).

The direct effect of reassortments on our results was moderate: between 67% and 91% of pairs of substitutions at significantly epistatic site pairs were not separated by any reassortment events (Table S1). In over 75% of all significantly epistatic site pairs, the majority of consecutive substitutions did not span any reassortment events.

### Enrichment of functional sites among epistatic sites

Next, we investigated whether sites that were implicated in inter-gene epistasis occurred preferentially in parts of the HA and NA proteins with known functional significance.

**Inter-gene versus intra-gene epistasis.** We compared the sets of sites involved in inter-gene epistasis to sets of sites involved in intra-gene epistasis, which we had identified previously [21]. The overlap between these two groups of sites was slightly higher than expected by chance for the H3 leading and the N2 trailing sites in the (H3,N2) gene pair (Table 2), but this difference was not significant after Bonferroni correction.

**Table 2.**
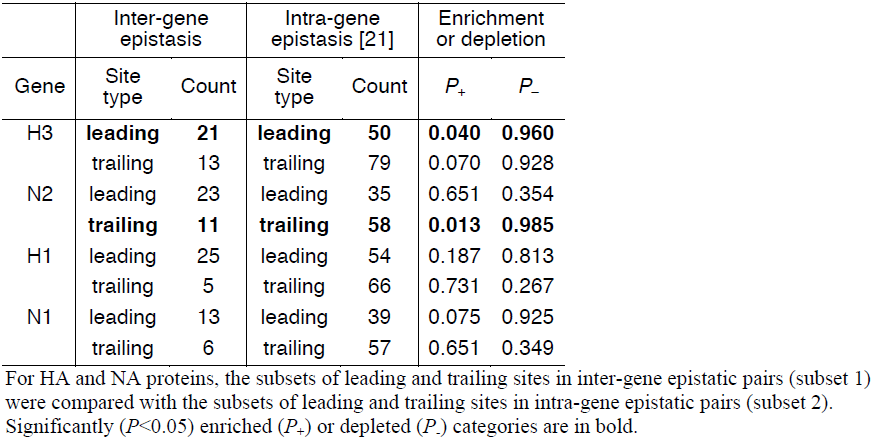
Comparisons of sets of sites evolving under inter-gene vs. intra-gene epistasis.

| Gene | Inter-gene epistasis |  | Intra-gene epistasis [21] |  | Enrichment or depletion |  |
| --- | --- | --- | --- | --- | --- | --- |
| | Site type | Count | Site type | Count | $P_+$ | $P_-$ |
| H3 | **leading** | **21** | **leading** | **50** | **0.040** | **0.960** |
|  | trailing | 13 | trailing | 79 | 0.070 | 0.928 |
| N2 | leading | 23 | leading | 35 | 0.651 | 0.354 |
|  | **trailing** | **11** | **trailing** | **58** | **0.013** | **0.985** |
| H1 | leading | 25 | leading | 54 | 0.187 | 0.813 |
|  | trailing | 5 | trailing | 66 | 0.731 | 0.267 |
| N1 | leading | 13 | leading | 39 | 0.075 | 0.925 |
|  | trailing | 6 | trailing | 57 | 0.651 | 0.349 |

For HA and NA proteins, the subsets of leading and trailing sites in inter-gene epistatic pairs (subset 1) were compared with the subsets of leading and trailing sites in intra-gene epistatic pairs (subset 2). Significantly ( $P < 0.05$ ) enriched ( $P_+$ ) or depleted ( $P_-$ ) categories are in bold.

**Epistatic sites and protein function.** The sites in HA identified as interacting with NA occurred in all parts of HA protein, with the majority of them located in known antigenic epitopes. This is expected because our method has more power to identify epistasis at sites that are more variable, and most of the variable sites in H3-HA1 are epitopic. Using a permutation-based approach (which controls for this bias), we found no statistically significant enrichment or depletion of epitopic sites, or sites under positive selection, among the epistatically interacting sites of HA or NA (Table 3). Trailing sites in H3 were slightly underrepresented among sites responsible for differences between antigenic clusters [34,35] and among sites that carry substitutions correlated with antigenic properties of isolates [36], as well as among glycosylation sites [37] (Table 3). Other than that, no obvious patterns in distribution of epistatically interacting sites in HA emerged.

**Table 3.** Comparisons of sets of sites evolving under inter-gene epistasis with sets of sites with known properties.

| Gene | Inter-gene epistasis |  | Category |  |  | Enrichment or depletion |  |
| --- | --- | --- | --- | --- | --- | --- | --- |
| | Site type | Count | Site type | Reference | Count | $P_+$ | $P_-$ |
| H3-HA1 | leading | 19 | **antigenic** | **[34]** | **44** | **0.023** | **0.975** |
|  |  |  | **antigenic** | **[36]** | **49** | **0.035** | **0.965** |
|  |  |  | antigenic | [35] | 7 | 0.255 | 0.745 |
|  |  |  | epitopic | [70,71] | 131 | 0.128 | 0.873 |
|  |  |  | glycosylation | [37] | 7 | 0.253 | 0.748 |
|  | trailing | 9 | **antigenic** | **[34]** | **44** | **0.993** | **0.005** |
|  |  |  | **antigenic** | **[36]** | **49** | **1.000** | **0.000** |
|  |  |  | **antigenic** | **[35]** | **7** | **0.990** | **0.008** |
|  |  |  | epitopic | [70,71] | 131 | 0.670 | 0.333 |
|  |  |  | **glycosylation** | **[37]** | **7** | **0.978** | **0.020** |
| H3-HA0 | leading | 21 | positively selected | - | 17 | 0.355 | 0.648 |
|  | trailing | 13 | positively selected | - | 17 | 0.945 | 0.053 |
| N2 | leading | 23 | epitopic | [71–73] | 45 | 0.620 | 0.378 |
|  |  |  | positively selected | - | 10 | 0.593 | 0.405 |
|  | trailing | 11 | epitopic | [71–73] | 45 | 0.158 | 0.840 |
|  |  |  | positively selected | - | 10 | 0.558 | 0.440 |
| H1-HA1 | leading | 21 | antigenic | [75] | 41 | 0.160 | 0.838 |
|  |  |  | epitopic | [20] | 32 | 0.133 | 0.865 |
|  |  |  | glycosylation | [76] | 11 | 0.528 | 0.470 |
|  | trailing | 5 | antigenic | [75] | 41 | 0.708 | 0.295 |
|  |  |  | epitopic | [20] | 32 | 0.138 | 0.863 |
|  |  |  | glycosylation | [76] | 11 | 0.958 | 0.040 |
| H1-HA0 | leading | 25 | positively selected | - | 6 | 0.588 | 0.415 |
|  | trailing | 5 | positively selected | - | 6 | 0.318 | 0.680 |
| N1 | leading | 13 | epitopic | [74] | 12 | 0.138 | 0.863 |
|  |  |  | **positively selected** | **-** | **2** | **0.993** | **0.005** |
|  |  |  | glycosylation | [76] | 13 | 0.713 | 0.285 |
|  | trailing | 6 | epitopic | [74] | 12 | 0.160 | 0.838 |
|  |  |  | positively selected | - | 2 | 0.253 | 0.745 |
|  |  |  | glycosylation | [76] | 13 | 0.130 | 0.868 |

For HA and NA proteins, the subsets of leading and trailing sites in inter-gene epistatic pairs (subset 1) were compared with the subsets of leading and trailing sites in a range of other categories (subset 2). Significantly ( $P < 0.05$ ) enriched ( $P_+$ ) or depleted ( $P_-$ ) categories are in bold.

Similarly, we found no significant enrichment of epitopic, positively selected, or glycosylation sites among sites in NA that are involved in inter-gene epistasis (Table 3). To better understand what types of sites in NA comprise the epistatic set, we searched the literature for evidence of functional consequences of mutations at sites that we identified. (Such a systematic analysis was impractical for HA because much less site-specific functional data is available for this protein.) We carried out this search for each of the 31 distinct sites in N2 (23 leading, 11 trailing sites, and 3 sites falling into both types), and for each of the 19 epistatic sites in N1 (13 leading and 6 trailing).

We found that inter-gene epistasis in both N1 and N2 may be related to NA catalytic activity and resistance to inhibitors (Table S2). Specifically, we found that N2 sites 370, 372, 401 and 432, which are homologous to the second sialic-binding site (hemadsorption site) of avian influenza viruses, are the leading sites in 58% (14/24) of the discovered epistatic pairs, including 13 pairs with the lowest *P*-values, and that they are trailing sites in 10% (3/29) of the discovered epistatic pairs (Table S2). Although the function of these sites in human influenza has not been directly demonstrated, it is thought that they affect catalytic efficiency of NA [38]. Among the remaining epistatic sites, leading sites 126 and 248 and trailing site 127 affect binding of NA inhibitors. In addition, two more leading sites, 215 and 332, although not shown to affect NA activity, were reported to often mutate in response to NA inhibitor treatment. Finally, two leading sites, 172 and 399, and one trailing site 263 were previously inferred to distinguish the reassortant H3N2 clades [32].

In N1, three leading sites 59, 386 and 388 (70, 390, and 392 in N2 numbering), and trailing site 434, undergo host-specific position-specific glycosylation, likely affecting enzymatic activity of NA [39]. Leading sites 6 and 14 and trailing site 15 are located in the transmembrane domain, which affects viral sialidase activity through its effect on NA tetramer assembly and transport to the membrane [40,41]. Leading site 149 and trailing sites 83, 275, 267 and 287 (274, 266, and 286 in N2 numbering) affect sialic acid binding; mutations at sites 267 and 275 were also shown to affect resistance to oseltamivir, including the mutation at site 275 which gave rise to the common oseltamivir-resistant H1N1 subtype. Finally, mutations at leading site 78, and trailing site 83, arise as compensatory in oseltamivir-resistant strains.

### Examples of epistatically interacting pairs of sites

Finally, we present several examples of implicated epistatic site pairs with biologically plausible explanations for the mechanism of their epistatic interactions.

**Leading site 78 in N1, trailing site 153 in H1.** We found a strong signal of epistasis between sites N1-78 and H1-153 (site 156 in H3 numbering; Table S3, Figure 2). Site N1-78 was also implicated in an intra-gene epistatic interaction with site N1-275 (site 274 in N2 numbering) [21], mutation at which causes oseltamivir resistance [42]. Substitutions at site N1-78 predated H275Y substitutions at least twice (Figure 2). Importantly, mutations at site 153 in H1 are known to be responsible for changes in receptor binding affinity [20], suggesting that a single mutation (N1-78) may precipitate further functionally important mutations in multiple genes.

**Figure 2.**
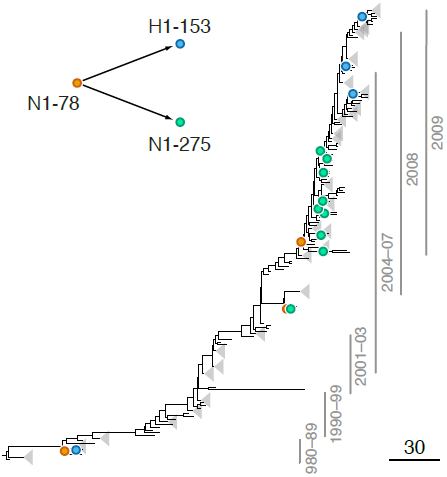
Example of putative inter-gene epistasis between sites N1-78 and H1-153. Substitutions at sites N1-78 and H1-153 are marked by orange and blue circles, respectively; substitutions at site N1-275 are marked by green circles. Site N1-275 was found to form a highly scoring inter-gene epistatic pair with site N1-78 in our previous study [21] (see text for details). Only substitutions that form consecutive pairs are shown (see Methods). Vertical bars show years in which the isolates where sampled. The inset shows the inferred directionality of epistatic interactions with arrows pointing from the leading to the trailing sites.

**Leading site 222 in H1, trailing sites 274 and 430 in N1.** We found a strong signal for epistatic interactions between site H1-222 (site 225 in H3 numbering) and sites N1-274 and N1-430 (Table S4, Figure 3), both of which, in turn, were implicated in intra-gene epistasis with site N1-275 [21]. Interestingly, mutations at site N1-430 modify activity of NA [43], while mutations at site H1-222, which is part of the receptor binding site, have been shown to compensate mutations in NA that confer resistance to NAI [44]. Since resistance to NAI depends on the balance between catalytic activities of HA and NA [23,45], mutations at these sites may be important for maintaining this balance.

**Figure 3.**
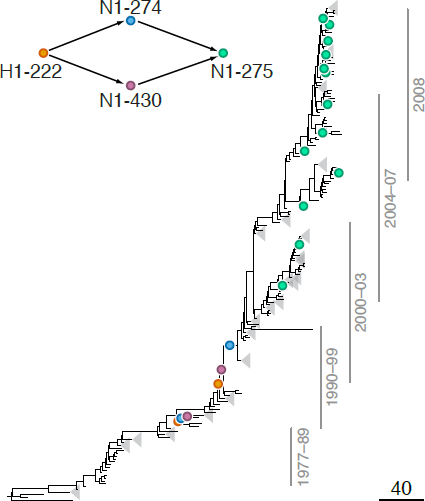
Example of putative inter-gene epistasis between sites H1-222 and N1-274. Substitutions at sites H1-222 and N1-274 are marked by orange and blue circles, respectively; substitutions at site N1-275 are marked by green circles. Site N1-275 was found to form a highly scoring inter-gene epistatic pair with site N1-274 in our previous study [21] (see text for details). Only substitutions that form consecutive pairs are shown (see Methods). The inset shows the inferred directionality of epistatic interactions with arrows pointing from the leading to the trailing site.

**Leading site 126 in N2, trailing sites 63 and 81 in H3.** We found a strong signal of epistasis between leading site N2-126 and trailing sites H3-63 and H3-81 (Figure 4). Site N2-126 frequently mutates in MDCK lines [46], and the observed substitution H126P is implicated to be important for the avian to human host shift of the H3N2 subtype [47]. Sites H3-63 and H3-81 are parts of known glycosylation motifs [37,48]. The loss of glycosylation site at position 81 in 1974 follows the gain of glycosylation site at position 63 in 1973 soon after beginning of H3N2 pandemic in 1968 [48], possibly in response to the H126P substitution in N2. Mutations at site H3-63 in three independent lineages created a new glycosylation site, while the old glycosylation site was concordantly lost either via a mutation at site H3-81 (twice) or a mutation at site H3-83 (once; Figure 4). Glycosylation of HA often masks epitopes [48,49] and loss of glycosylation at site 81 may also affect receptor binding [50]. We speculate that adaptation to the new host occurred via a change in receptor-binding activity of NA that in turn precipitated compensatory mutations in HA glycosylation patterns.

**Figure 4.**
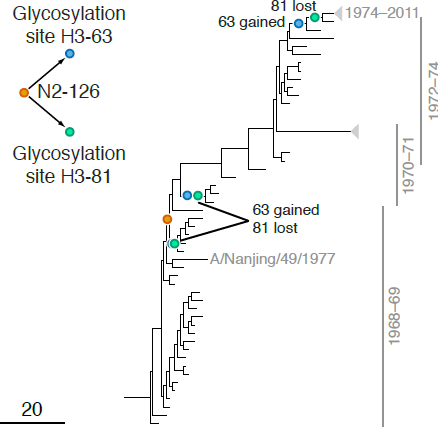
Example of putative inter-gene epistasis between sites N2-126 and H3-63 and H3-81. Substitutions at sites N2-126 and glycosylation motifs starting at sites H3-63 and H3-81 are marked by orange, blue and green circles, respectively. All three substitutions affecting the H3-63 glycosylation motif occurred at site H3-63 and created a new glycosylation site. Two substitutions affecting the H3-81 glycosylation motif occurred at site H3-81 and one occurred at site H3-83, but all of them destroyed an existing glycosylation motif. Vertical bars show years in which the isolates where sampled. The inset shows the inferred directionality of epistatic interactions with arrows pointing from the leading to the trailing site.

## Discussion

Here we developed a phylogeny-based method for detecting positive epistasis between mutations at sites that are incompletely linked. This approach provides the first systematic procedure for identifying such genetic interactions from sequence data sampled over time. We demonstrated the power of this method by applying it to data from human influenza A virus where we found dozens of putative epistatic interactions between sites in the surface proteins HA and NA. Several of the most significant pairs of sites implicated by this statistical procedure have known biological functions that provide a plausible mechanistic basis for the observed patterns of coordinated molecular evolution.

While powerful, our method of detecting epistasis between incompletely linked sites suffers from two important limitations. First, it relies on our ability to accurately infer phylogenies, detect reassortment events and map substitutions from one phylogeny onto another. Accurate detection of reassortments, especially between very closely related taxa, is difficult [28,31], and the mapping of substitutions is inherently ambiguous. Although the precise effect of such reconstruction errors on the performance of our method is unclear, we expect that they would not lead to an inflation of the epistasis signal, at least as long as the number of reassortment events is small. The second limitation is inherent to the problem of detecting epistasis from temporal substitution data, and is discussed in detail in a previous study using such techniques [21]. The problem is that our method (as well as any method utilizing the same data) will identify sites as trailing in epistatic pairs if substitutions at these sites are temporally clustered for any reason – including reasons that are not caused by epistatic interactions *per se*. Site N1-275, mutations at which confer resistance to oseltamivir, is a likely example. Most substitution at this site occurred in the span of just 3 years between 2006 and 2009. Based on such highly non-random distribution of substitution events on the phylogeny, our method implicates this site in many inter- as well as intra-gene epistatic interactions with sites that experienced substitutions shortly prior to this period [21]. While the experimental evidence confirms that some of these sites genuinely interact in determining viral fitness [19,51], some of the other inferred interactions may be spurious.

Keeping these caveats in mind, we turn to the interpretation of our observation of epistasis between mutations in the HA and NA. The numbers of discovered epistatic site pairs where the leading mutation occurs in NA or in HA are similar. Thus, substitutions in HA facilitated by prior substitutions in NA and substitutions in NA facilitated by prior substitutions in HA appear to be equally common. The evolution of these two segments of the human influenza virus is therefore tightly coordinated.

What is the molecular basis for such coordinated evolution? We searched for enrichment of various properties among epistatically interacting sites. In HA, we found no enrichment of epistatic sites in any of the characterized functional categories. Sites implicated in epistatic interactions are neither sites responsible for antigenic shifts nor sites evolving under positive selection. In fact, it is somewhat surprising that the detected epistatic sites are not particularly rapidly evolving, despite the fact that our method has more power to detect epistasis at sites with more substitutions [21]. Thus, epistatic sites in HA comprise a novel potentially interesting set of sites in this protein.

We also found no enrichment of positional or functional categories in epistatic sites in NA. This lack of clear pattern is consistent with experimental data and implies that genetic interactions occur through a wide range of mechanisms, and that the sites involved in them are hard to predict a priori [50,51]. However, we observed that many epistatic sites in NA are involved in NAI resistance, modulation of NA activity, or both. Why do the sites affecting these traits interact with HA? Some of the observed interactions (e.g., site N1-78 (Figure 2), and sites N1-274 and N1-430 (Figure 3)) could be directly attributed to the requirement to balance the activities of HA and NA to maintain viral fitness, especially in the presence of NAI [45]. Other interactions may affect this balance indirectly. For example, sites in the signal peptide of HA appear to occasionally interact with sites in the transmembrane domain of NA, e.g., site H1-16 forms a putatively epistatic pair with site N1-15 (Table S6). These types of mutations likely affect the efficiency of membrane localization of the respective surface proteins [40], and mutations in the transmembrance domain may also influence NA activity through their effect on tetramer assembly [41].

Some of the putatively epistatic site pairs that we detected have been experimentally confirmed. For example, a number of substitutions in HA of H1N1 closely predated the 2007 spread of the H275Y (274 in N2 numbering) oseltamivir resistance substitutions in NA. Recently, 7 of these HA sites were experimentally tested for interactions [51]. These experiments showed that HA that carries the derived residues at all seven sites is well adapted to both the ancestral H275 (sensitive) and the derived Y275 (resistant) variant of NA. At the same time, three out of seven reconstructed reversions in HA (at sites 82, 141 and 189) had large fitness defects in the context of the derived NA variant, implying that substitutions at these sites compensated for the H275Y substitution in NA [51]. Remarkably, all three of these HA sites form high-ranking pairs in our analysis with the site 275 in NA (Table S6, sites 99, 157, 205 in our numbering).

More generally, our results suggest that the evolution of a protein depends strongly on its genomic context, with a substantial number of adaptive mutations representing responses to mutations that previously occurred in other proteins. Such evolutionary coupling between different proteins has also been observed in several experimental systems [13,15,23,52–54]. However, estimating the fraction of substitutions that are driven by direct adaptation to the external environment versus by selection to balance or compensate the effects of prior substitutions elsewhere in the genome remains an important open problem.

## Materials and Methods

### Sequences

We downloaded all complete human H3N2 influenza A isolates (N=2,205) available on 27 October 2011 and all complete human seasonal H1N1 influenza A isolates (N=1,180) available on 12 November 2012 from the flu database [55]. The amino acid sequences were aligned using MUSCLE [56,57], and the alignments were reverse translated using PAL2NAL [58]. Genotypes containing truncated sequences or long stretches of unidentified nucleotides were discarded. The 3 genotypes of H3N2 subtype carrying indels were discarded. We also discarded all genotypes of H1N1 that were sampled prior to 1936 because they had large (15-16 amino acids) gaps between amino acid positions 42 and 77 in the NA protein. In all sequences, the alignment columns with gaps in more than 10% of all sequences were excluded from further consideration; in the remaining alignment columns, gaps were substituted with the consensus nucleotide.

Four isolates of H1N1 subtype (A/New Jersey/1976, A/Wisconsin/301/1976, A/Iowa/CEID23/2005, A/Switzerland/5165/2010) were discarded as swine-origin influenza virus (SOIV) [59–61]. Three isolates of H3N2 subtype (A/Ontario/RV123/2005, A/Ontario/1252/2007 and A/Indiana/08/2011) were discarded as SOIV triple reassortants [62].

Many of the genotypes had NA genes with identical nucleotide sequences; among each such set of genotypes, we only retained one random genotype. This reduced our sample to 1,376 isolates for H3N2 subtype, and 745 isolates for H1N1 subtypes.

For HA and NA proteins of H1N1, the numbering scheme used through the text is relative to the proteins of the A/AA/Huston/1945 isolate, unless stated otherwise.

### Inferring the temporal order of substitutions in two reassorting segments

We asked whether a substitution at a particular site in HA segment facilitates a subsequent substitution at a particular site in NA segment, or vice versa. To address this, we need to reconstruct the phylogenetic trees for each of the two segments, infer the position of reassortments on these trees, and establish the temporal order of substitutions in different segments relative to each other. We achieve this goal in three steps, which are described in detail below. Briefly, in the first step, based on topological incongruencies between the phylogenetic trees of individual segments, we identify the so-called reassortment sets, i.e., sets of taxa that are likely descendants of reassortant viruses. In the second step, we reconstruct the so-called constrained phylogenies of the segments, i.e., phylogenies that are topologically identical everywhere except for branches that correspond to reassortment events. This allows us to map, in the third step, the substitutions that occur on branches of one phylogeny to the branches of another phylogeny.

**Inferring the reassortment and the “trunk” sets.** We used GiRaF [31] to identify sets of taxa that are descendant to reassortment events. To reduce the computational burden associated with this step, we first clustered isolates with nucleotide identity exceeding 99.5% across the concatenated HA-NA sequence using CD-HIT [63], and retained for the GiRaF analysis one random sequence from each cluster, for a total of 225 H3N2 and 169 H1N1 isolates.

GiRaF takes as input the sets of phylogenetic trees sampled from their posterior distributions for each segment. We obtained 1000 such trees per segment using MrBayes [64] with the GTR+I+Γ model, 2 million iterations, sampling one tree every 2000 iterations. The output of GiRaF is a collection of taxon sets each of which consists of descendants of a likely reassortment event. Because GiRaF attempts to infer nested reassortments and because of phylogenetic noise, these sets are generally overlapping, i.e., the same taxon may be included into multiple sets. However, to infer subtrees with topologies unaffected by reassortments, we need non-overlapping sets of taxa each descendant to the same past reassortment event (or the same series of such events). To construct such non-overlapping sets, we sorted the GiRaF sets according to the fraction of taxa shared with other sets, from high to low. All taxa in the highest-ranking set were then considered as one set of reassortants. We then excluded these taxa from all lower-ranking sets, resorted the remaining GiRaF sets, and repeated the procedure. Thus, for example, if GiRaF set 1 was fully nested within a larger GiRaF set 2, we inferred two non-overlapping sets of reassortants: those of set 1, and those of set 2 excluding those of set 1. A GiRaF set not overlapping any other GiRaF sets always produced a set of reassortants of its own. By this procedure, each taxon was included either into a unique reassortment set (denoted by the most recent reassortment event), or into the set of non-reassortant taxa which we refer to as the “trunk” set. We then ascribed the isolates removed in the clustering step to the same set as their representative cluster sequence.

**Reconstructing constrained phylogenies.** Given *N* sets of taxa (*N*–1 reassortment sets and one trunk set), we reconstruct two complete phylogenies (one per segment) that differ by exactly *N*–1 rooted subtree prune-regraft (rSPR) operations corresponding to *N*–1 reassortment events. We call such phylogenies “constrained”. To assemble constrained phylogenies, we start by reconstructing two standard maximum likelihood phylogenies (one per segment) using PhyML [65] (model GTR+I+Γ) and rooting these phylogenies with the oldest isolate as the outgroup (A/Albany/18/1968 for H3N2, and A/Henry/1936 for H1N1). We use these phylogenies as templates for reconstructing constrained phylogenies.

Next, for each reassortment or trunk set of taxa, we reconstruct an unrooted phylogenetic subtree from the alignment of concatenated HA and NA sequences by maximum likelihood using PhyML [65] (model GTR+I+Γ). To root each such subtree, we compare the locations of the most recent common ancestors (MRCAs) of this set of taxa on two template trees. In the absence of phylogenetic noise, MRCAs on both segments would be identical, in which case the root of the concatenate-based subtree would be placed unambiguously. However, in general, MRCAs based on different template phylogenies are different. We therefore place the root of the concatenate-based subtree in such a way that its position is most similar to both alternative MRCA positions according to a trade-off function described in Text S1. As a result of this procedure, we obtain *N*–1 reassortment rooted subtrees and one trunk rooted subtree.

We then assemble these subtrees into two complete constrained phylogenies (one per segment) that differ by exactly *N*–1 rooted subtree prune-regraft (rSPR) operations as follows. In the absence of noise, i.e., if reassortments were the only source of differences between the two template phylogenies, each reassortment set would be either mono- or paraphyletic on each template phylogeny. Each reassortment subtree could then be unambiguously grafted into a unique branch of another (paraphyletic) subtree, in exact accordance with the template tree. However, some reassortment sets are polyphyletic on the template trees, making the grafting procedure ambiguous. Our algorithm resolves such ambiguities on the basis of a tradeoff between two criteria: maximizing topological similarity between the constrained and the template phylogeny for each segment, and minimizing the length of the resulting constrained phylogeny (see Text S1 for details).

The result of this assembly is a pair of phylogenetic trees, one tree per segment, that differ from each other by *N–*1 rSPR operations, as desired. Once the topologies of the constrained phylogenies are reconstructed, we optimize their branch lengths and infer ancestral sequences using HyPhy with the nucleotide REV+Rate Het. model [66].

**Establishing temporal order of events on the phylogeny.** Our goal is to detect substitutions in one segment that occurred after substitutions in another segment. If we analyze substitutions in segment 1 that occurred after substitutions in segment 2, we say that “segment 2 forms the genetic background for segment 1”, or that “segment 2 is in the background” and “segment 1 is in the foreground”.

To study such substitutions in segment 1, we map substitutions in the background segment (segment 2) onto the foreground-segment phylogeny. Since the topologies of each reassortment and trunk subtree are identical for both segments by construction (see above), each branch of the background tree maps to a unique branch of the foreground tree (we call such branches “unambiguous”), with the exception of branches that are involved in the rSPR operations (we call such branches “ambiguous”). Thus, substitutions in the background segment that occur on unambiguous branches map onto unique branches of the foreground phylogeny. Consider a toy example shown in Figure 1. Node *b* forms a single reassortment set, and the remaining nodes form the trunk set. Correspondingly, the constrained phylogenies shown for the two segments differ by a single rSPR operation involving the branch leading to node *b*. Therefore, branches *gb*, *ec*, and *ed* in the segment 2 phylogeny map to branches *fb*, *ec*, *ed* of the segment 1 phylogeny, respectively, and substitution *iv* in segment 2 unambiguously occurs on branch *fb* of the segment 1 phylogeny (Figure 1B).

Now consider ambiguous branches, such as branches *rg*, *ga*, and *re* in the segment 2 phylogeny in Figure 1A. Each rSPR corresponding to each reassortment event removes one node (prune operation), thus merging a pair of successive branches, and adds one node (regraft operation), thus splitting a branch. Therefore, each reassortment event results in one branch of the background phylogeny corresponding to a pair of branches of the foreground phylogeny (1-to-2 map) and another pair of branches of the background phylogeny corresponding to another branch of the foreground phylogeny (2-to-1 map). In Figure 1, a pair of branches *rg* and *ga* of the segment 2 phylogeny corresponds to branch *ra* of the segment 1 phylogeny, and branch *re* of the segment 2 phylogeny corresponds to the pair of branches *rf* and *fe* of the segment 1 phylogeny. In 2-to-1 maps, all substitutions that occur on either of the two branches in the background segment are unambiguously mapped onto a single branch of the foreground segment phylogeny. In Figure 1, substitution *ii* in segment 2 maps unambiguously onto branch *ra* of the segment 1 phylogeny. The situation is more difficult in the 1-to-2 maps, where each substitution that occurs on such ambiguous branch in the background segment could map onto either one of the two branches in the foreground segment. In Figure 1, substitution *iii* in segment 2 could occur either on branch *rf* or on branch *fe* of the segment 1 phylogeny. We resolve this ambiguity by placing all such background substitutions onto the distal branch of the foreground phylogeny (e.g., in Figure 1, substitution *iii* is placed on branch *fe*). This choice minimizes the number of consecutive potentially epistatic substitution pairs that can form between the background and the foreground sites (see below).

Finally, each reassortment event maps onto the branch of the foreground segment phylogeny that leads to the most recent common ancestor of the corresponding reassortment subtree. We refer to such branches as “reassortment-carrying branches”, or RCBs. We signify the occurrence of a reassortment event by adding a “virtual” node on the RCB and placing all foreground substitutions that occur on the RCB after the virtual node. Thus, we make a simplifying assumption that the reassortment event precedes all substitutions on the RCB. For example, in Figure 1B, reassortment (virtual node *h*) precedes substitutions *i* and *iv* on branch *fb* of segment 1 phylogeny. When viewed as an event on the foreground phylogeny, each reassortment event is equivalent to an instantaneous replacement of the background segment sequence present in the parent node of the foreground RCB with the background segment sequence present at the parental node of corresponding RCB on the background segment phylogeny. In the example shown in Figure 1B, the reassortment event replaces segment 2 which carries no mutations (present at node *f*) with a segment 2 descendent from node *g* that has mutation *ii*. Thus, this reassortment event is equivalent to the occurrence of substitution *ii* in the background segment. This “virtual” substitution is placed on the virtual branch *fh*. This procedure of mapping background substitution onto the foreground phylogeny guarantees that the order of background substitutions is preserved.

All procedures for construction of constrained gene trees were implemented in C++ with bio++ package [67,68]. Mapping of substitutions was implemented in Perl and used Bio::Phylo package [69].

### Inferring positive epistasis

**Epistasis statistic.** To infer positive epistasis between substitutions at two sites mapped onto the same phylogeny, we employ the method previously described in [21]. Briefly, for each pair of sites (*i,j*), we first identify the set *S*_*ij*_ of all consecutive substitutions pairs, i.e. such substitution pairs where a substitution at site *i* is on the line of descent of a substitution at site *j* with no other substitutions at either site occurring in between. We then compute the epistasis statistic for this pair which in the simplest case is given by

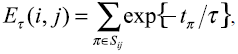

where the summation is taken over all consecutive substitution pairs, *t*_π_ is the time (measured in synonymous substitutions) between the substitutions in the pair π, and τ is the time-scale parameter which we choose to be equal to the average time 〈*t*_π_〉 averaged over all substitutions at all site pairs (Table 1). For a more general expression of the epistasis statistic see [21]. The epistasis statistic for site pair (*i,j*) is large when (a) the set of consecutive substitution pairs is large and (b) when the substitutions at the trailing site quickly follow the substitutions at the leading site. Thus, site pairs with an unusually high epistasis statistic likely evolve under positive epistasis.

**Identifying epistatic site pairs and computing the false discovery rate (FDR).** In order to identify site pairs with unusually high values of the epistasis statistic, we obtained the null distribution for the epistasis statistic at all site pairs simultaneously by randomly reshuffling substitutions at all sites among all branches of the phylogeny. Our permutation procedure conserves the number of substitutions at each site and on each branch thus controlling for possible biases introduced by differences in variability among sites and by the heterogeneity of substitutions on the phylogeny [21]. We carried out 10,000 permutations for each analysis. Using the resulting null distributions for each site pair, we obtained the list of site pairs that were significant at any given nominal *P*-value (observed positives, OP). To estimate the number of false positives (FP) that we expect to find at a given nominal *P*-value, we selected 400 out of 10,000 permutations as fake data sets and calculated the number of significant site pairs in each of these fake data sets at that *P*-value. Thus, we obtained the null distribution of the number of significant pairs for each nominal *P*-value, which allowed us to estimate the expected number of FP (EFP) under the null hypothesis as well as the *P*-value for the number of OP. The FDR is then given by the ratio EFP/OP.

### Testing for functional enrichment among sites involved in inter-gene epistasis

Lists of epitopic sites of HA were taken from [70,71] for H3N2 and from [20] for H1N1. Lists of epitopic sites of NA were taken from [71–73] for H3N2, and from [74] for H1N1. Sites involved in intra-gene epistasis in HA and NA of H3N2 and H1N1 were taken from [21]. Sites that may carry substitutions changing the antigenic properties of isolates were taken from [34–36] for H3, and from [75] for H1. Glycosylation sites were taken from [37] for H3-HA1, and from [76] for H1-HA1 and H1-NA. Rapidly evolving sites in H1, N1, H3 and N2 were inferred by HyPhy IFEL method [77] (p-value < 0.05) using Datamonkey web service [78,79] (http://www.datamonkey.org).

To test whether a particular set of sites *S* is enriched or depleted among the top-ranking epistatic leading sites compared to the random expectation, we used the following procedure. First, we defined for each site its leading *P*-value as the lowest nominal *P*-value among all site pairs with this site as leading. We then defined the leading test statistic as the difference between the medians of site's leading *P*-value for sites in *S* and for sites not in *S*. The null distribution of this test statistic was determined from the 400 fake datasets generated from the no-epistasis null hypothesis as explained above. An analogous procedure was used to find enrichment among the top-ranking epistatic trailing sites.

## Supplementary Materials

**Text S1**. Building constrained phylogenies.

**Table S1**. Distribution of locations of consecutive substitution pairs relative to reassortment events.

**Table S2**. Properties of epistatic sites in N1 and N2 neuraminidases.

**Table S3**. Putatively epistatic site pairs identified in the (H3,N2) gene pair at nominal *P*-value threshold 0.05. Notations for the location of consecutive substitutions relative to reassortment events. First letter denotes whether the leading and the trailing substitutions are located on the same (S) or different (D) reassortment subtrees. Second and third letters denote the branch types on which the leading and the trailing substitutions are located: R = reassortment-carrying branch; V = virtual branch; I = any other internal branch.

**Table S4**. Putatively epistatic site pairs identified in the (N2,H3) gene pair at nominal *P*-value threshold 0.05. Notations as in Table S3.

**Table S5**. Putatively epistatic site pairs identified in the (H1,N1) gene pair at nominal *P*-value threshold 0.05. Notations as in Table S3.

**Table S6**. Putatively epistatic site pairs identified in the (N1,H1) gene pair at nominal *P*-value threshold 0.005. Notations as in Table S3.

**Figure S1**. The number of observed (orange line) and expected (black line) discovered epistatic pairs at each nominal *P*-value threshold in different gene pairs. One (two) asterisks denote cases when the observed number exceeds the expectation according to the permutation test at significance level 0.05 (0.01). See Materials and Methods for details.

